# Ubiquitinated protein levels remain high during ageing in *C. elegans*

**DOI:** 10.1101/2023.11.07.566011

**Authors:** Sandrine E. Daigle, Savannah Doucet, Louis R. Lapierre

**Affiliations:** Centre de Médecine de Précision du Nouveau Brunswick, 27 rue Providence, Moncton, New Brunswick, E1C 8X3, Canada; Département de chimie et biochimie, Université de Moncton, 18 Antonine Maillet, Moncton, New Brunswick, E1A 3E9, Canada; Department of Molecular Biology, Cell Biology and Biochemistry, Brown University, 185 Meeting St., Providence, Rhode Island 02912, USA

## Abstract

A wealth of literature in different model organisms have shown that proteostasis declines with age, which results in the accumulation of aggregated proteins that generally carry polyubiquitin chains. In Koyuncu et al. (Nature, 2021), the authors use a relatively mild homogenization approach to analyze total polyubiquitinated protein levels longitudinally in *C. elegans* and found that the overall level of polyubiquitinated proteins decreases in wild-type nematodes. Specifically, the substantial presence of polyubiquitinated proteins in early adulthood markedly decreases at day 10 and 15 of adulthood. This finding represents the central basis of their study where they subsequently suggest that age-dependent decrease in protein polyubiquitination in wild-type animals is due to enhanced deubiquitination. Here, we find that the mild homogenization of *C. elegans* used in Koyuncu *et al*. fails to properly extract all *C. elegans* proteins, because it largely omits relatively insoluble proteins in the pellet. When we perform complete homogenization of wild-type nematode proteins using SDS, sonication and heat, we unequivocally observe that polyubiquitinated protein levels do not decrease with age in *C. elegans*. Overall, the levels of polyubiquitinated proteins remains relatively constant post-reproduction. Therefore, our findings invalidate the main conclusion of Koyuncu et al. that the ubiquitinated proteome is rewired during ageing and demonstrate that *C. elegans* harbor polyubiquitinated proteins throughout their lifespan.

## INTRODUCTION

It is well documented that the proteostasis network in *C. elegans* begins to fail relatively quickly during their adulthood (1, 2), which leads to loss of protein solubility (3) and progressive accumulation of proteins in aggregates (4-6). Aggregated proteins tend to generally be polyubiquitinated, and they may or may not be amenable to proteasomal degradation, depending on their size, the availability and accessibility of chaperones and disaggregases, and the overall activity of the proteasome (7, 8). Therefore, proper inquiries into protein ubiquitination requires the analysis of both soluble and insoluble proteins (9). Protein aggregates also impair the activity of the proteasome, thereby increasing the likelihood of further accumulation of unprocessed, unstable and aggregation-prone polyubiquitinated proteins (10). Notably, significant protein aggregation has been observed in middle aged wild-type *C. elegans* (4-6).

The foundation of the Koyuncu et al. (11) study lays largely on the observation in Figure 1g (and Supplementary Figure S2a) that wild-type animals, unlike long-lived *daf-2* and *eat-2* animals, display an unexpected and substantial decrease in polyubiquitinated proteins with age (clearly visible at and beyond Day 10 of adulthood), using protein immunoblotting. The authors later suggest that deubiquitinase may be involved in generating those changes in older wild-type animals. In order to understand the mechanism by which polyubiquitinated protein levels (or any other types of proteins for that matter) are modulated with age, it is imperative that the entirety of present proteins in nematodes be analyzed. However, the authors use a mechanical homogenization approach in cold conditions with limited amount of detergent (1% NP-40 and 0.25% sodium deoxycholate), which is not particularly well suited to fully solubilize proteins in nematodes. Potentially, a significant portion of polyubiquitinated proteins may be missed in their study, rendering the lysis and analysis incomplete, and consequently the interpretation problematic and likely erroneous. We therefore asked: Are polyubiquitinated protein levels in wild-type animals decreasing when complete protein solubilization is performed? Here, by using a protocol that completely solubilizes proteins in *C. elegans*, we unambiguously demonstrate that the level of polyubiquitinated proteins ubiquitination does not decrease during ageing. Therefore, we refute the premise of “rewiring” of the ubiquitin proteome with ageing in *C. elegans* as suggested in Koyuncu *et al*. We conclude that the omission of the insoluble protein pellet during the lysis of *C. elegans* across different approaches in the publication of Koyuncu *et al*. render their analyses incomplete and misleading. Altogether, our findings contradict the observations in Koyuncu *et al*. on the reduction of protein ubiquitination with age in wild-type *C. elegans* and put into question the importance and relevance of the subsequent findings in Koyuncu et al. related to the role of deubiquitinase in protein ubiquitination, proteostasis and ageing in *C. elegans*.

**Figure 1.**
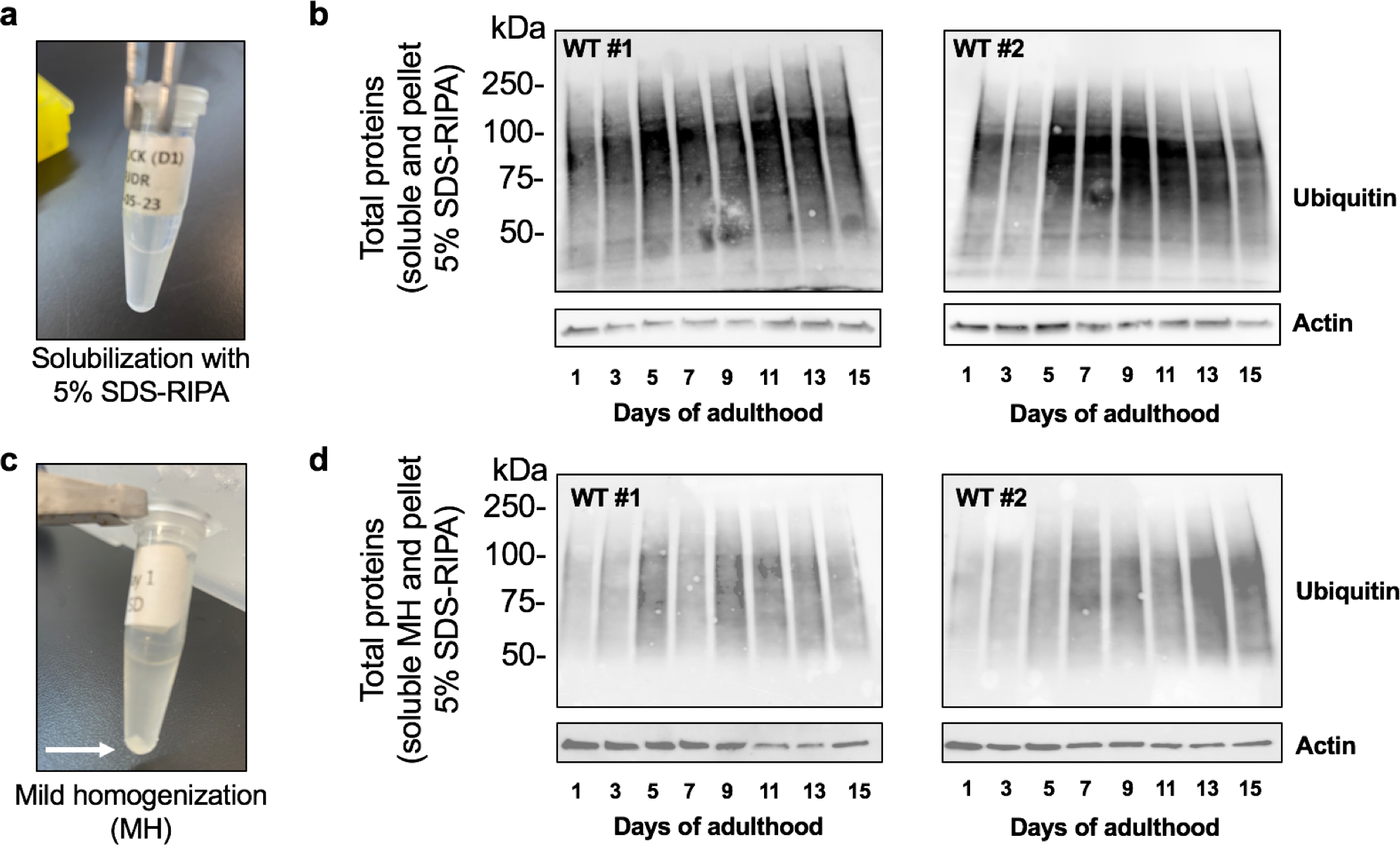
Polyubiquitinated protein levels remain elevated throughout lifespan in *C. elegans*. **a**. Visualization of homogenate post-centrifugation after the first solubilization with 5% SDS-RIPA buffer (Day 1 sample). **b**. Immunoblotting of ubiquitinated proteins and actin in wild-type nematodes (#1, Kenyon lab, #2, Tuck lab) from Day 1 to Day 15 of adulthood. **c**. Visualization of homogenate post-centrifugation after the first solubilization with the mild homogenization (MH) buffer from Koyuncu et al. (Day 1 sample – white arrow highlights noticeable pellet). **d**. Visualization of homogenate post-centrifugation after the first solubilization with the mild homogenization buffer from Koyuncu et al.

## RESULTS

In order to convincingly determine whether protein polyubiquitination levels changes during ageing in *C. elegans*, we opted to carry out complete protein homogenization in a 5% SDS-containing RIPA buffer (see methods), which promotes solubilization of all proteins, including insoluble protein aggregates. For each nematode collection (approximately 500), complete homogenization required two rounds of solubilization, whereas the nematodes were solubilized with 5% SDS-RIPA, heated and sonicated, and the small remaining pellet was re-solubilized accordingly. Approximatively 500 worms were collected every two days from Day 1 to Day 15 of adulthood. Before collecting, we picked out the dead worms to ensure that only live worms were studied (it is unclear whether Koyuncu *et al*. actually removed dead worms in their analyses). Removing dead nematodes is particularly important beyond Day 9 of adulthood when a significant portion of wild-type animals start dying, as the mean lifespan of wild-type animals at room temperature is approximately 15 days. As in Koyuncu *et al*., the sterilizing agent floxuridine (FUDR) was used to sterilize the animals and facilitate longitudinal nematode collection. N2 wild-type animals originating from the Kenyon laboratory (N2 #1, likely same origin than Koyuncu *et al*. via A. Dillin lab at Salk/UC Berkeley) were grown on NG plate seeded with OP50 *E. coli* and sterilized with FUDR (as described in Koyuncu *et al*.). An alternative N2 strain from another nematode laboratory (12) (N2 #2) was used to control for potential lab-to-lab wild-type variations. Complete solubilization was performed in two steps whereas remaining insoluble pellet from the initial solubilization was resolubilized (**Figure 1a**). Polyubiquitinated proteins from combined soluble and solubilized pellet fractions (total) were visualized by immunoblotting using the same anti-ubiquitin antibody as in Koyuncu *et al*. (see Methods). In both wild-type N2 strains, total solubilization of proteins with 5% SDS-RIPA clearly demonstrated that the levels of polyubiquitinated proteins does not decrease during ageing in *C. elegans* (**Figure 1b**). Using the homogenization protocol from Koyuncu *et al*., we found that a substantial pellet remained after lysis (**Figure 1c**). Notably, we were able to completely solubilize this pellet with 5%SDS-RIPA, heat and sonication. Combining the soluble fraction from this mild homogenization with the corresponding fully solubilized pellet fraction similarly demonstrated that polyubiquitinated protein levels do not decrease with age in wild-type animals (**Figure 1d**). Altogether, our data clearly shows that the levels of polyubiquitinated proteins in N2 wild-type animals do not decrease with age. Instead, they remain relatively constant, suggesting that nematodes maintain a steady burden of polyubiquitinated proteins throughout their life. Since Koyuncu *et al*. utilized an incomplete lysis protocol for the entirety of their study, our study seriously challenges their conclusion on the rewiring of the ubiquitin proteome during ageing.

## METHODS

### Nematode synchronization and collection

Wild-type animals were grown on OP50 *E. coli* and synchronized by bleaching for experiments. Eggs were seeded on agar plates containing OP50 *E. Coli* and sterilized with FUDR as described in Koyuncu *et al*. Worms were transferred onto fresh plates every 5 days. Approximately 500 worms were collected per time points for each solubilization method.

### Protein solubilization for immunoblotting

To completely solubilize proteins, we used a RIPA solution (50mM Tris-HCl, 150mM NaCl, 1mM EDTA, Roche Complete Protease Inhibitors) containing 5% SDS followed by heating for 5 minutes at 95°C and sonication for 20 seconds (QSonica Q55, 30% input). Insoluble proteins were pelleted by centrifugation for 2 minutes at 12000rpm. This first solubilization step (250μl of 5% SDS-RIPA) solubilized over 90% of total proteins, and a second step of solubilization (250μl of 5% SDS-RIPA) successfully solubilized the small pellet. Ubiquitinated proteins from combined (i.e. total) were visualized by immunoblotting. Mild homogenization (MH) was also performed using the buffer (250μl) and method described in Koyuncu et al. (11) and insoluble proteins were pelleted by centrifugation for 2 minutes at 12000rpm. Subsequently, these larger pellets (compared to the 5% SDS-RIPA method) were completely solubilized with 5% SDS-RIPA, heat and sonication as described above. Polyubiquitinated proteins were visualized by immunoblotting for ubiquitin (Millipore Sigma 05-994, clone P4D1-A11) and actin was used as loading control (Millipore Sigma MAB1501R, clone C4).

## ACKNOWLEDGEMENTS

We would like to thank Dr. Brad Morrison, Sophie Landry and Karine Blais for their feedback. We would like to thank the Malene Hansen laboratory for the N2 wild-type strains used in this study. SED and SD performed the immunoblotting and edited the manuscript. LRL designed and performed the experiments, and wrote the manuscript. This work was funded by an NSERC Discovery Grant and a Research Chair in Precision Medicine (J.-Louis Lévesque Foundation – NB Research) to LRL.

## CONFLICT OF INTEREST

None declared.

### DATA AVAILABILITY STATEMENT

All data discussed in this manuscript are presented in Figure 1.

